# What’s next? Predictable visual sequences are encoded with anticipatory biases and reduced neural responses

**DOI:** 10.1101/2025.10.16.682933

**Authors:** Songyun Bai, Maëlan Q. Menétrey, David Pascucci

## Abstract

Objects in motion follow predictable trajectories that the brain can easily anticipate. We investigated the underlying neural mechanisms, focusing on a form of representational momentum (RM), whereby the final state of a rotating object is misperceived along its future trajectory. Participants viewed two simultaneous streams of oriented gratings, one rotating predictably and the other random, and were instructed to attend to one of the two. Using EEG steady-state visual evoked potentials (SSVEPs) and decoding analysis, we found that attending to rotational sequences led to reduced neural response amplitude and systematic anticipatory shifts in the neural representation of orientation, relative to random sequences. These anticipatory biases emerged at early post-stimulus latencies and mirrored behavioral signatures of RM. Our findings suggest that the brain leverages internalized dynamics to form stable anticipatory representations with reduced neural activity strength.

## Introduction

Most events in our visual world unfold according to familiar and predictable physical principles—a glass pushed off a desk falls, and a ball thrown against a wall bounces back. These regularities allow the visual system to anticipate the trajectory and future states of objects (Kubricht et al., 2017; Rosen, 2012), often leading to systematic anticipatory biases. A striking example is representational momentum (RM), in which the perceived final state of a dynamic or transforming object—such as one undergoing rotational motion—is misreported in the direction of its implied trajectory (Bertamini, 2002; Freyd & Finke, 1984; Hubbard, 2005; Pascucci & Plomp, 2021). RM has been widely interpreted as a behavioral signature of predictive mechanisms in perception, especially for stimulus dimensions that involve continuous change and are frequently encountered in natural contexts (Erlhagen, 2003; Hafri et al., 2022; Hubbard, 1995, 2005, 2010; Jancke & Erlhagen, 2010; Kerzel, 2003; Müsseler et al., 2002).

Despite extensive behavioral evidence for RM and its central relevance to predictive processing, the underlying neural mechanisms remain poorly understood. In the broader literature on predictive perception, one of the most consistently reported neural signatures is *expectation suppression*—a reduction in neural responses to predictable stimuli (Alink et al., 2010; Han et al., 2019; Richter et al., 2022; Walsh & McGovern, 2018). These effects are often interpreted within the framework of predictive coding, which posits that top-down predictions generated in higher cortical areas suppress sensory signals in early visual regions when incoming input is highly predictable (Alink et al., 2010; Kok & de Lange, 2015; Rao & Ballard, 1999; Trapp et al., 2021). However, most empirical studies of predictive coding have focused on static or discrete visual inputs, often employing explicit cues or block-wise manipulations to induce expectations (e.g., Kok et al., 2013, 2017; Aitken et al., 2020). These paradigms, while informative, do not capture the temporally extended, naturalistic dynamics of real-world perception—precisely the kind of contexts in which anticipatory biases like RM emerge. As such, it remains unclear whether expectation suppression or related predictive coding mechanisms apply to perceptual extrapolation in continuous dynamic scenarios.

There is also debate about the processing stage at which RM arises. Some evidence points to the involvement of low-level mechanisms, such as motion extrapolation signals already in the retina (Berry et al., 1999). At the cortical level, motion-sensitive regions like MT/V5 respond not only to actual movement but also to static images that imply motion (Kourtzi & Kanwisher, 2000; Senior et al., 2000; but see Rao et al., 2004). Additionally, recent work suggests that the visual system can pre-activate stimulus-specific sensory templates in anticipation of dynamic, unfolding input, reflecting predictive processing across multiple stages (Blom et al., 2020; de Vries & Wurm, 2023; Ekman et al., 2017). However, RM-like effects have also been linked to higher-order regions, including prefrontal cortex, implicating the involvement of more cognitive and mnemonic processes (Rao et al., 2004). In general, behavioral studies indicate that RM unfolds rapidly, within 250–300 ms after stimulus offset (Freyd & Johnson, 1987), but it is unclear whether this reflects a post-perceptual bias during working memory encoding or a genuinely predictive modulation—i.e., a shift in neural representations that already occurs during initial processing.

A further unresolved issue concerns the role of attention. While some findings suggest attention is necessary for anticipatory motion extrapolation (Kerzel, 2003), others report enhanced RM when attention is diverted, implying an automatic, possibly pre-attentive origin (Hayes & Freyd, 2002). Supporting this, the magnitude of RM correlates with neural indices of automatic deviance detection such as the visual mismatch negativity (Kimura, 2018), indicating that forward perceptual biases may arise early in sensory processing even without explicit top-down modulation.

Thus, although RM provides a valuable window into anticipatory mechanisms in perception, fundamental questions remain regarding its influence on neural response amplitude, the processing stage where stimulus representations begin to shift forward, and the role of attention.

In the present study, we address these questions using an EEG design combining steady-state visual evoked potentials (SSVEPs) with inverted encoding models (IEMs). We tracked the neural encoding of sequences of oriented stimuli that either followed a predictable rotational structure or varied randomly. This design allowed us to test whether temporal regularities in the input modulate not only the amplitude of visual responses but also the content of neural representations—specifically, whether they exhibit anticipatory forward shifts consistent with RM. Importantly, we manipulated attention by asking participants to attend either to the structured or the random sequence, presented concurrently. This enabled us to assess whether predictive shifts in neural representations depend on attentional engagement or instead arise automatically from exposure to temporal regularities.

## Results

We recorded EEG while nineteen participants viewed two lateralized sequences of flickering gratings (Figure□1a; see Methods). Each sequence lasted 5□seconds and flickered at 5□Hz, yielding 25 gratings per stream, each lasting 200ms. Grating orientations changed on every flicker, in steps of 20° or integer multiples thereof. On each trial, participants were cued to attend to one of the two streams while maintaining central fixation. After the sequence offset, they judged whether a centrally presented probe matched the orientation of the final grating in the attended stream.

In the rotational (RTT) sequence, orientations followed a predictable clockwise progression with fixed 20° steps. In the random (RDM) sequence, orientation changes also occurred in multiples of 20° but varied unpredictably in both direction and magnitude. For clarity, we refer to these as “rotational” and “random” sequences, although both involved stepwise orientation rotations. The two types of sequence were randomly assigned to the left or right visual field on each trial. Throughout the manuscript, we refer to RTT condition as the condition in which participants were cued to attend to the RTT sequence, and RDM condition when they were cued to attend to the RDM sequence.

This design allowed us to assess how temporal structure and attention jointly modulate perceptual performance, SSVEP amplitude, and the fidelity of orientation-specific neural representations across the stimulus stream.

### Representational momentum in perceptual reports

We evaluated the presence of the typical forward displacement bias in RM by analyzing the proportion of ‘same’ reports—i.e., the proportion of times the probe orientation was reported as the same as the last attended orientation, as a function of the temporal structure of attended sequence (RTT vs. RDM). This analysis was conducted using a 2-way repeated-measures ANOVA with factors: (1) the relative orientation difference between the probe and the last grating in the attended sequence (-4°, 0°, or +4°) and (2) the condition (attend to RTT vs. RDM).

As shown in Figure 1c, the results of the ANOVA revealed no main effect of condition (F(1,18) < .001, *p* = .982). However, there was a significant main effect of the relative orientation difference (F(2,36) = 9.964, *p* < .01), and a significant interaction between condition and orientation difference (F(2,36) = 6.761, *p* < .01).

**Figure 1:**
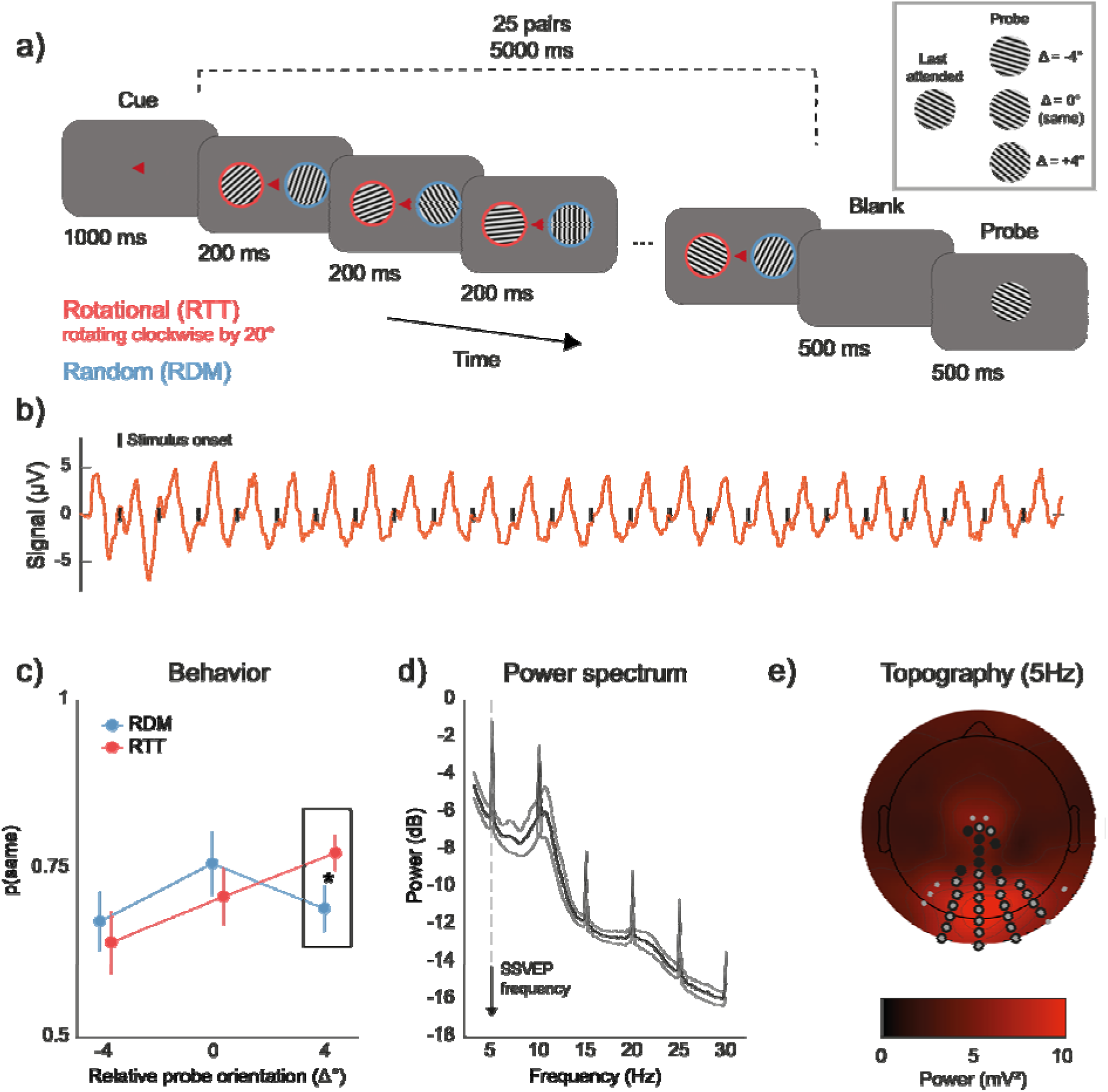
a) Experimental design. Each trial began with a small red triangle cueing the side that participants were instructed to attend to. One second later, two flickering gratings with rotational (RTT) and random (RDM) sequences were simultaneously displayed in both visual fields. At the end of the trial, participants indicated whether the probe orientation matched the last orientation in the attended sequence. b) Steady-state visual evoked potential (SSVEP). The typical periodic EEG response evoked by the flickering stimulus is shown at an example electrode A25 (approximately Iz in the standard 10-20 system) for an example subject. c) Representational momentum (RM) in behavioral reports. In the RTT condition (in red), participants tended more to report the probe as similar to the last attended orientation when it was shifted by 4°, indicating that the last grating in the sequence was perceived as shifted forward along the future trajectory of rotation. No such bias was observed in the RTT condition (in blue). Error bars are 95% CI, corrected for within-subject design (Cousineau, 2005). d) SSVEP effects on the power spectrum. The power spectrum across all electrodes revealed a clear peak at the SSVEP frequency (5 Hz) and its harmonics (e.g., 10 Hz). Shaded areas are 95% CI. e) Topography of power distribution at 5 Hz. The topographical distribution of 5 Hz power, averaged across subjects, is shown. Electrodes with power exceeding the 75th percentile at both 5 Hz and 10 Hz are highlighted in gray and black, respectively, and were subsequently used for decoding analysis (see Methods).

A post-hoc t-test indicated that this interaction was driven by a higher proportion of ‘same’ responses in the RTT condition (when participants attended to the rotational sequence) when the relative orientation difference was +4° (t(18) = 3.00, *p* = .008, Cohen’s d = 0.688). That is, in the RTT condition, the orientation of the probe stimulus had to be shifted forward by 4° to be perceived as identical to the last attended stimulus.

Furthermore, in the RTT condition, the proportion of ‘same’ responses at +4° was significantly higher than at 0° (t(18) = 3.03, *p* = .007, Cohen’s d = 0.696) and at -4° (t(18) = 5.45, *p* < .001, Cohen’s d = 1.251). No such pattern was observed in the RDM condition: participants were most likely to respond ‘same’ when the probe matched the last stimulus exactly (0°), with significantly fewer ‘same’ responses at -4° (t(18) = 2.58, *p* = .019, Cohen’s d = 0.591) and +4° (t(18) = 3.00, *p* = .008, Cohen’s d = 0.555). Although our design tested only a limited range of orientation differences compared to prior RM studies (e.g., Pascucci & Plomp, 2021), these results indicate that perceptual judgments in the RTT condition were systematically biased forward along the rotation trajectory—a hallmark of representational momentum (Hubbard, 2005).

### Attentional modulation of SSVEPs is attenuated by rotational sequences

The power spectral density of the SSVEPs showed clear increases at the stimulus frequency (5 Hz) and its first harmonic (10 Hz), with spectral peaks most prominent over occipito-parieto-central electrodes (Figure 1d and 1e).

We then investigated how SSVEP responses were modulated by attention and by the temporal structure of the sequences. Consistent with prior findings that attention enhances SSVEPs contralateral to the attended visual field (Norcia et al., 2015), we assessed lateralized effects separately for the RDM and RTT conditions.

In the RDM condition, time-frequency analysis revealed a marked increase in 5 Hz power contralateral to the attended side (paired t-test, left vs. right, cluster-based permutation statistics at the electrodes and time points level, *p* < .01; Figure 2a & 2b, left panel). In the RTT condition, however, no significant lateralized effects were observed at any electrode or time point (Figure 2a & 2b, right panel).

**Figure 2:**
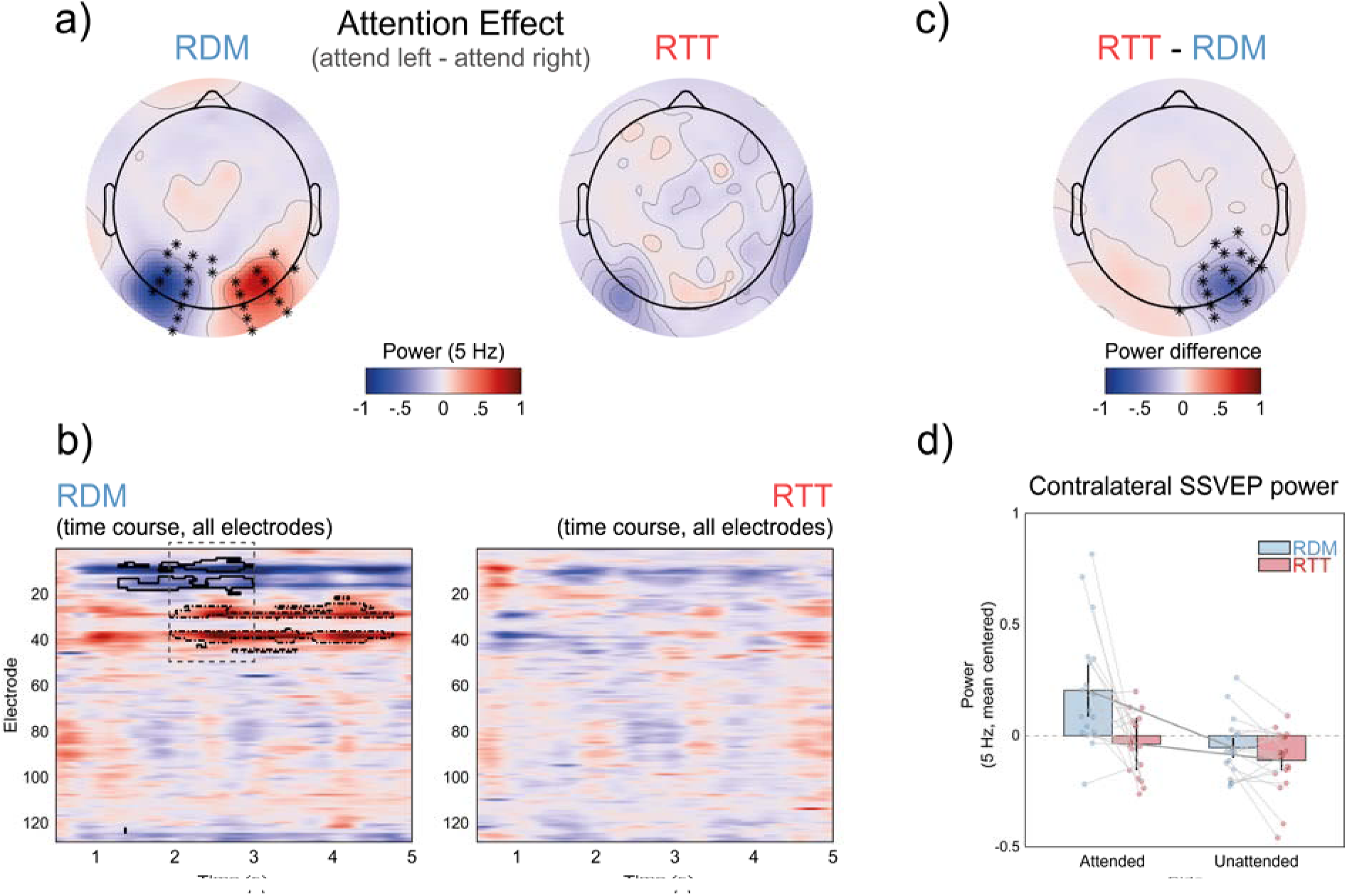
Effects of attending to rotational or random visual sequences on SSVEP. a) Topographies of attention effects. The averaged power difference at 5 Hz is shown for left-attended minus right-attended sequences in both random (RDM) and rotational (RTT) conditions. Significant electrodes are marked with asterisks. Notably, no electrodes showed significant effects in the RTT condition. b) Time-frequency analysis. The two panels illustrate the 5 Hz power difference (left-attended minus right-attended) for the two conditions. Significant electrode-time clusters are outlined in black, indicating lateralized effects of spatial attention. The dashed rectangle marks the time window where both left- and right-attended stimuli induced significant effects, corresponding to the topographies shown in (a). c) Topography of condition differences. The map highlights the cluster showing reduced lateralization of the SSVEP attention effect in the RTT condition compared to the RDM condition. This difference was calculated after collapsing across the attended side and flipping electrode positions to align with the right-attended condition. d) Contralateral SSVEP power. The SSVEP power at the contralateral significant cluster identified in (c), averaged across the whole trial, provides a measure of lateralization at the stimulation frequency across the four conditions (RDM in blue, RTT in red). Error bars represent 95% confidence intervals, and individual participant values are shown as points. For visualization purposes, each participant’s mean across conditions was subtracted from their data.

To further assess this difference, we computed a lateralization index of SSVEP activity (see Methods) and compared it between conditions. The results showed a significant reduction in contralateral SSVEP activity in the RTT condition (Figure 2c; paired t-test, left vs. right, cluster-based permutation statistics at the electrodes and time points level, *p* < .01). A two-way repeated-measures ANOVA performed on the power at 5 Hz at contralateral sites (Figure 2d), averaged over the whole epoch, revealed a main effect of attention (attend left vs. right, F(1,18) = 9.48, *p* < .01) and of temporal structure (RTT vs. RDM, F(1,18) = 10.083, *p* < .01) as well as an interaction between the two (F(1,18) = 5.815, *p* = .027), driven by a significant lateralization effect only evident when attending to the RDM sequence (attended RDM vs unattended RDM, t(18) = 3.087, *p* < .01, attended RTT vs unattended RTT t(18) = 1.783, *p* = .092).

Together, these results suggest that while attention typically enhances SSVEP responses contralateral to the attended hemifield, this effect is markedly attenuated in the presence of structured rotational dynamics.

### Anticipatory neural encoding of rotational sequences

We then assessed the neural representation of individual stimulus orientations in the RDM and RTT conditions—i.e., when participants attended to rotational or random sequences. To this aim, we used an inverted encoding model (IEM) (Sprague et al., 2018) to reconstruct orientation-selective population responses at the electrode sites showing the strongest SSVEP responses (Figure 1e; 5 Hz and 10 Hz). The IEM was trained on trials from the RDM condition (see Methods) and then tested on both RDM and RTT trials, using a cross-validation procedure to decode the orientation of each grating at each time point, from 200 ms before stimulus onset to 500 ms after stimulus offset.

The IEM results are shown in Figure 3a. Decoding orientations in the RDM condition revealed two significant clusters of time points: an early cluster from 72 to 180 ms post-stimulus, and a later cluster from 240 to 312 ms (both *ps* < .05). The decoder trained on the RDM condition also generalized to the RTT condition, successfully reconstructing the orientation tuning function between 80 to 268 ms post-stimulus (*p* < .05; Figure 3a).

**Figure 3.**
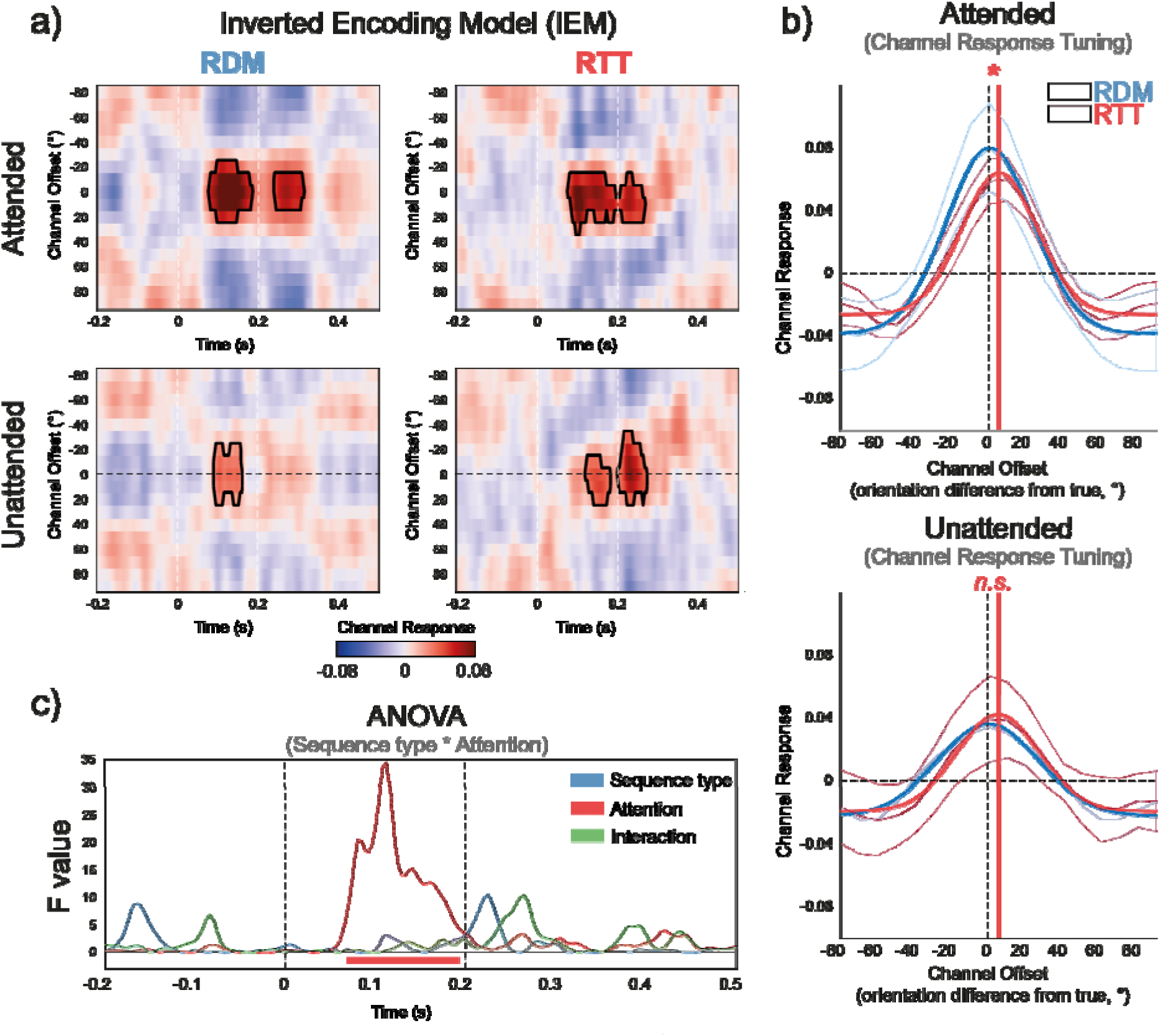
Inverted Encoding Model. a) Reconstructed tuning function over time. The reconstructed tuning function for model orientation channels, realigned to the true stimulus orientation (0° by convention), is shown across time for the four conditions. The figure is organized by sequence type (RDM and RTT, in separate columns) and attention (Attended vs. Unattended, in separate rows). Solid black lines indicate time clusters where decoding accuracy was significant (*p* < .05). b) Tuning functions within significant clusters. Reconstructed tuning functions averaged over significant time clusters. These tuning functions were fitted with Gaussian models, with results for the RDM and RTT conditions shown in blue and red, respectively. The tuning curves reflect the strength and precision of stimulus orientation representation. In the RDM condition, the function is centered on the actual stimulus orientation (0°), indicated by the black dashed vertical line. In the attended RTT condition, the function is shifted 5° clockwise (red solid vertical line), reflecting an anticipatory bias towards the future orientation. No significant bias is observed in the unattended RTT condition, though a similar trend is present (bottom plot in b). c) Effects of attention and sequence type on decoding scores. F-values from repeated-measures ANOVA on decoding scores are shown, evaluating the main effects of sequence type, attention, and their interaction. Significant effects are indicated by the horizontal line at the bottom (*p* < .05). A significant main effect of attention is observed at the time of significant decoding clusters (indicated by the red horizontal line), with no effect of sequence type or interaction.

By averaging the channel tuning functions (CTFs) within the first significant cluster for both conditions and fitting a Gaussian function, we found a significant forward shift in the peak decoded orientation in the RTT condition. Specifically, the peak orientation was shifted by approximately 5° toward the future trajectory of rotation (RTT peak decoded orientation vs. zero, *p* < .01, jackknife statistics, see Methods; Figure 3b). This anticipatory bias appeared early in the post-stimulus period, emerging within the first significant cluster before 100 ms, suggesting modulation during early stages of visual processing.

We also performed the decoding analysis for unattended sequences. Significant decoding was observed in both unattended RDM and RTT conditions. While the unattended RTT condition exhibited a numerically similar forward bias to the attended RTT condition, this effect did not reach statistical significance (Figure 3b). No significant differences in the width of the Gaussian fits were found across conditions.

Finally, we assessed how attention and sequence type influenced the fidelity of neural representations. Decoding accuracy scores (see Methods) from -200 to 500 ms relative to stimulus onset were analyzed with a time-resolved two-way repeated-measures ANOVA including factors of attention (attended vs. unattended), sequence type (RTT vs. RDM), and their interaction. As shown in Figure 3c, attention significantly enhanced decoding fidelity within the time window of significant orientation decoding (from 68ms to 192ms after stimuli onset, time-resolved cluster-based statistics, *p <* .01). No main effect of sequence type or interaction was observed.

### Efficient encoding of rotational sequences with reduced neural activation

Our findings thus far indicate two key results: (1) attending to the RTT sequence, compared to the RDM sequence, leads to reduced SSVEP lateralization; and (2) despite a forward bias in orientation decoding for RTT stimuli, the reliability of neural representations remains comparable across conditions. Together, these findings suggest that rotational sequences may be encoded with reduced neural response strength while preserving representational fidelity.

To further evaluate this hypothesis, we examined a more general index of EEG signal magnitude: the Global Field Power (GFP; see Methods). As shown in Figure 4, GFP was consistently higher in the RDM condition than in the RTT condition (Figure 4a, 4c). Despite this, decoding accuracy for individual grating orientations remained comparable between the two conditions (Figure 4b, 4c). This dissociation between overall signal strength and representational fidelity supports the conclusion that rotational dynamics are encoded more efficiently—i.e., with reduced neural resources but without a loss in the quality of stimulus representation.

**Figure 4.**
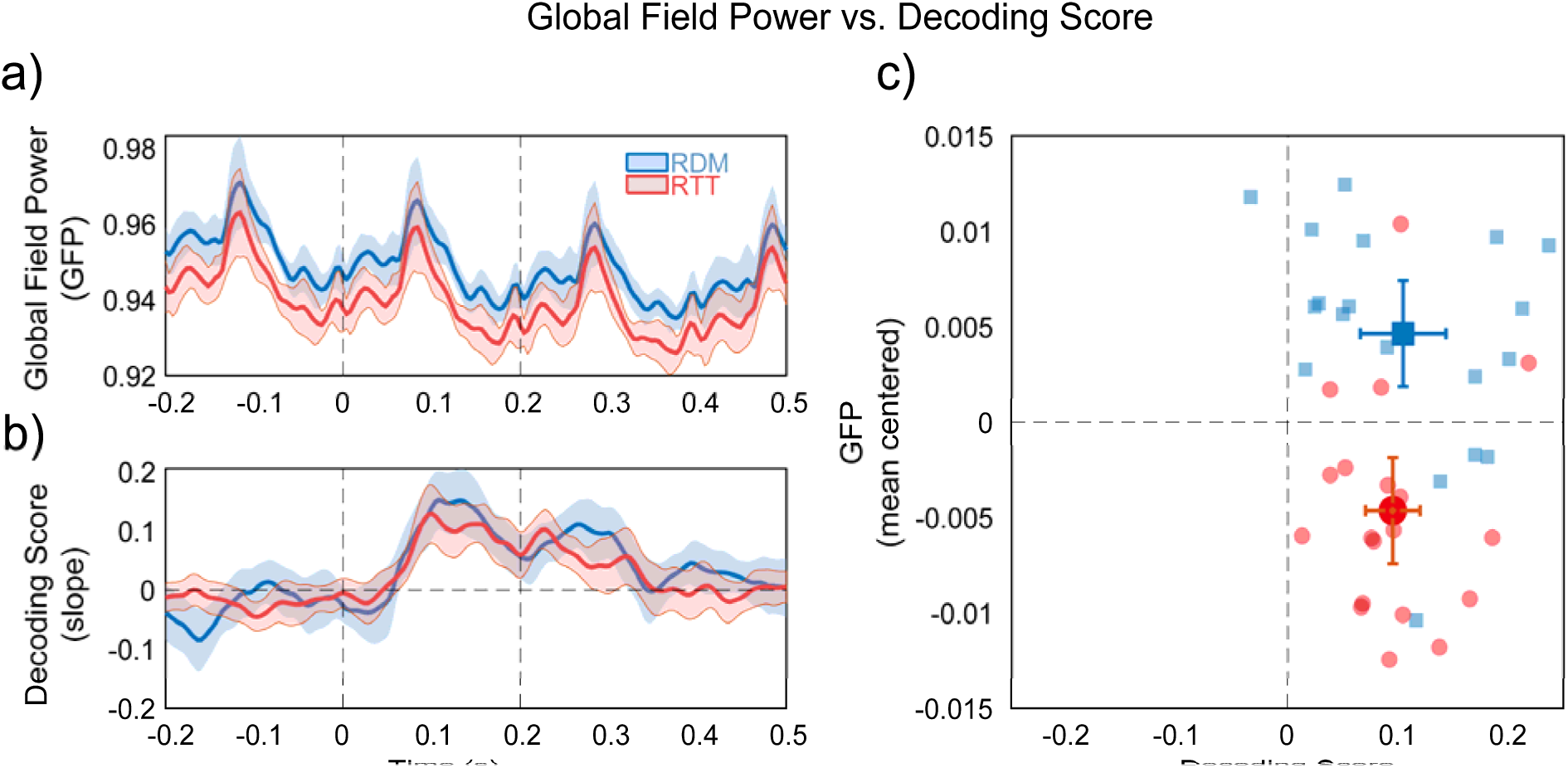
Global field power (GFP) computed as the standard deviation of signals across the scalp at each time point, for the RDM (in blue) and RTT (in red) conditions, within a time window from -200 to 500 ms relative to the current stimulus. b) Decoding scores for the two conditions in the same time window as in a. c) A plot showing the relationship between decoding scores (x-axis) and GFP (y-axis), with the overall GFP mean per participant subtracted to facilitate visual comparison. The same color coding is applied for each condition as in a and b. Individual data points represent individual participants. Shaded areas in a and b, as well as error bars in c, represent 95% CI. Dashed vertical lines in a and b mark the onset and offset of the current stimulus.

## Discussion

In this study, we investigated how attention to structured visual sequences influences both the amplitude of neural responses and the content of neural representations, as assessed via SSVEPs and EEG-based decoding of stimulus orientation.

Our results demonstrate that attending to rotational sequences leads to a reduction in overall EEG amplitude, including attenuated lateralized SSVEP responses—a neural signature typically enhanced by spatial attention. Despite this reduction in signal amplitude, orientation decoding accuracy remained comparable between structured (RTT) and random (RDM) sequences. Decoded orientation representations in the RTT condition exhibited a systematic anticipatory bias in the direction of rotation. This forward shift emerged early in the post-stimulus time window and closely aligned with the behavioral RM effect.

Below, we discuss the implications of these findings for current theories and debates on predictive processing, consider potential underlying mechanisms, and outline the limitations of our study.

### Rotational sequences are reliably encoded with reduced neural activity

One of the main findings was the reduction in lateralized SSVEP power and GFP during attention to rotational (RTT) compared to random (RDM) sequences. Since rotational sequences were highly predictable, this effect could be interpreted within the framework of expectation suppression (Alink et al., 2010; Han et al., 2019; Richter et al., 2022; Walsh & McGovern, 2018) and predictive coding (Friston, 2005; Rao & Ballard, 1999).

As mentioned, predictive coding posits that when sensory input is highly predictable, top-down signals suppress bottom-up responses, which may explain the attenuated neural activity in the RTT condition. This attenuation could, in principle, also coincide with a reduction of decodable stimulus information (Richter et al., 2018): because internal models already account for expected input, there may be little unpredicted sensory data to encode, and hence less stimulus information to decode from neural activity. However, we found comparable decoding accuracy across conditions, with decoders trained on RDM trials generalizing well to RTT trials. This suggests that stimulus information was preserved across conditions, speaking against a strong predictive coding account in which both neural responses and stimulus representations are silenced by top-down predictions.

Other views propose that top-down predictions may suppress prediction error signals while still preserving and refining stimulus representations—for instance, by sharpening neural tuning around the encoded stimulus features or by reducing internal noise (Kok, Jehee, et al., 2012; Walsh & McGovern, 2018). In these views, reduced EEG amplitude could reflect more efficient or selective encoding, without a loss of decodable information. However, our data do not support such sharpening mechanisms either. We found no differences in decoding accuracy or in the width of the reconstructed channel tuning functions between conditions. This suggests that neither a sharpening nor an attenuation of orientation-selective representations occurred—at least as indexed by EEG-based decoding in the context of RM.

Thus, while our results are partly consistent with core assumptions of predictive coding, they point to a more nuanced pattern.

### Neural representations of rotational sequences exhibit forward bias

To our knowledge, this study provides the first direct evidence that stimuli implying rotational motion are encoded with anticipatory forward shifts toward their future states. This finding extends previous work demonstrating that the brain can preactivate future positions of regularly moving objects, effectively compensating for neural processing delays (Blom et al., 2020; Ekman et al., 2017; Hogendoorn, 2020; Hogendoorn & Burkitt, 2018; Johnson et al., 2023). Here, we observed a forward shift in the decoded stimulus feature—orientation rather than spatial position—within an early temporal window beginning approximately 80 ms post-stimulus onset, specifically when participants attended to rotational sequences. This anticipatory bias was already qualitatively apparent at the first decoding peak (∼80–100 ms, Figure 3b), suggesting that predictive representations arise rapidly, likely within early visual cortex (Ester et al., 2020).

These findings speak to debates about the locus of RM, which has been attributed to both perceptual and mnemonic processes (Brouwer et al., 2004; Hubbard, 2005; Pascucci & Plomp, 2021; Reed & Vinson, 1996). Our data indicate that the forward shift occurs swiftly after stimulus onset, supporting the interpretation that RM encompasses an early perceptual component rather than solely reflecting post-perceptual memory biases.

### Attention enhances stimulus-related neural content and forward biases

Our findings are also relevant to the role of attention in processing regular sequences of visual events. Prior studies have emphasized the joint influence of attention and expectation, often assuming they exert opposing effects on neural activity: attention enhances, while expectation suppresses responses (Alink & Blank, 2021; Garrido et al., 2018; Kok, Rahnev, et al., 2012). This opposition model would predict the strongest neural responses when attention is directed to unpredictable stimuli—as in our attended RDM condition—and reduced responses when predictable stimuli are unattended, as in our unattended RTT condition (Figure 2d). However, this view primarily focuses on overall neural response magnitude. Our findings demonstrate a dissociation between neural activity strength and the fidelity of neural representations: while rotational sequences led to reduced EEG amplitude (SSVEP and GFP), stimulus decoding remained robust across conditions. Moreover, attentional engagement enhanced decodability in both conditions, and the magnitude of anticipatory forward biases in the RTT condition, even in the presence of reduced activity.

This pattern is not easily reconciled with models based solely on neural activity strength. Instead, it may reflect an interaction between attention and anticipatory processing that selectively enhances task-relevant content without increasing overall activity.

### Candidate neural mechanisms

Several mechanisms may explain the presence of anticipatory bias despite an overall reduction in neural activity during structured rotational sequences. First, early elements in the sequence might trigger automatic neural cascades that preactivate future orientations, decreasing the need for continuous sensory tracking (Ekman et al., 2017, 2023; Johnson et al., 2023). Second, the visual system may rely on low-dimensional latent representations that encode motion direction and predicted positions, rather than processing each frame independently (Hénaff et al., 2021). Third, short-term adaptation may reduce responses to recently seen orientations while enhancing sensitivity to slightly forward-tilted ones, thus producing a shift aligned with the motion trajectory (Ceylan & Pascucci, 2023; Clifford et al., 2007; Pascucci et al., 2019; Pascucci & Plomp, 2021). Fourth, within the biased competition framework (Desimone & Duncan, 1995), attention resolves competition between concurrently presented stimuli by biasing processing toward behaviorally relevant features. In our task, one could hypothesize that this bias operates more efficiently in structured sequences, where predictive mechanisms pre-activate expected orientations (Ekman et al., 2017), reducing the need for bottom-up amplification while preserving representational precision.

While our data cannot distinguish between these explanations, they converge on a consistent conclusion: structured rotational sequences elicit anticipatory neural encoding that is both efficient and systematically biased in the direction of motion, emerging from the earliest stages of visual processing.

### Limitations

A key limitation of our study is that the behavioral investigation was primarily confirmatory, designed around the minimal conditions necessary to detect the presence of representational momentum (RM). Specifically, the sparse sampling of orientation differences between the probe and the last attended stimulus limited our ability to fully characterize the shape and peak of the behavioral RM bias. This, in turn, prevented meaningful comparisons and correlations between EEG decoding results and behavioral measures. Nonetheless, the behavioral effects we observed are consistent with classic RM findings and with more recent studies using similar designs (Gottwald et al., 2015), supporting the broad conclusion that our manipulation induced biases characteristic of RM.

Additionally, we employed only a single rotation direction—clockwise. This approach was chosen to induce systematic expectations across trials and to maximize the number of epochs per condition for EEG analysis. We see no theoretical reason to expect different results with counterclockwise rotations, but this remains to be empirically verified.

Another limitation arises from the use of IEMs. The models rely on assumed shapes for channel tuning functions and were trained exclusively on the RDM condition. This may have complicated direct comparisons of changes in tuning width—and thus representational precision—between the RDM and RTT conditions. Lastly, the current design could not disambiguate whether the lateralized SSVEP responses in the RTT condition were also partly driven by inhibitory activity suppressing the processing of predictable but unattended sequences on the ipsilateral side (Zhao et al., 2013), which remains a question for future research.

## Conclusion

Our study provides evidence that the brain processes regularly evolving sequences of visual stimuli with reduced neural activity and anticipatory representations. While the precise underlying mechanisms remain to be fully understood, we propose that the brain employs such efficient strategies when encoding naturally evolving sequences, such as rotational dynamics, to minimize neural resources while preserving reliable representations of stimulus trajectories.

## Methods

### Participants

Nineteen healthy participants (8 women, 11 men; mean age 21.3 years, SD 1.2 years) were recruited from École Polytechnique Fédérale de Lausanne (EPFL) and the University of Lausanne (UNIL). Participants received monetary compensation (30 CHF/h) for their participation. All participants had normal or corrected-to-normal vision, assessed through the Freiburg acuity test (threshold for inclusion: >1; 2), and were naive to the purpose of the study. The protocol was approved by the local ethics committee and adhered to the Declaration of Helsinki. Before the experiment, participants provided written informed consent and were informed of their right to withdraw at any time.

### Apparatus

The study was conducted in the Laboratory of Psychophysics (LPSY) at EPFL. Participants were seated in a dimly lit, electromagnetically shielded room, 100 cm from an ASUS VG248QE LCD monitor (1920 × 1080 pixels, 24.5-inch screen, 144 Hz refresh rate). Stimuli were generated and presented using custom scripts in MATLAB (R2022b, The MathWorks, Natick, USA) and the Psychophysics Toolbox (Brainard, 1997).

### Stimuli and procedure

We combined steady-state visual evoked potentials (SSVEP) with an experiment assessing representational momentum (RM). On each trial, participants viewed two simultaneously presented sequences of grating stimuli—one in the left and one in the right visual hemifield (Figure 1a). Each grating subtended 15° of visual angle, had a spatial frequency of 0.5° and a Michelson contrast of 0.1, and was centered ±8° horizontally from the center of the screen.

Grating pairs were presented for 200 ms each with no interstimulus interval, yielding a stimulation frequency of 5□Hz. This frequency was selected for three main reasons. First, prior work suggests that anticipatory processes may be mediated by oscillations in the theta band (∼4–7□Hz), making ∼5□Hz a functionally relevant frequency for probing prediction-related mechanisms (Han & VanRullen, 2017). Second, this rate avoids overlap with the alpha band (∼10□Hz), where endogenous EEG activity is strong and could reduce the signal-to-noise ratio of SSVEPs. Third, higher stimulation frequencies would require faster transitions between orientations, deviating from typical temporal parameters used in RM paradigms, which often involve stimulus durations longer than 200□ms and include interleaved blanks (Hubbard, 2005; Pascucci & Plomp, 2021).

In one of the sequences—the RTT (rotational) condition—grating orientations followed a regular, clockwise progression in 20° steps. In the RDM (random) condition, orientations were drawn from the same set but presented in a randomized order, with the constraint that no two consecutive gratings formed a 20° transition. This removed any local rotational structure while keeping the overall orientation statistics matched across conditions. The design, with the two conditions, follows previous work (Kelly & Freyd, 1987; Kerzel, 2002; Pascucci & Plomp, 2021), and was specifically tailored to assess how regularity and predictability influence neural responses and the decoded representation of each stimulus relative to a matched but unpredictable control.

Each trial began with the presentation of a small red isosceles triangle at the center of the screen, serving as a spatial attention cue. The triangle’s base measured 0.6°, and its apex—also 0.6° away—pointed either left or right, indicating the hemifield to which participants should direct their attention. Participants were instructed to maintain central fixation while covertly attending to the cued sequence. The triangle remained visible throughout the 5-second stimulation period (25 grating pairs).

Importantly, the attention cue determined the condition for that trial: when the cue pointed toward the regularly rotating sequence, the trial was classified as RTT condition; when it pointed to the randomly ordered sequence, the trial was classified as RDM condition.

After a 500□ms blank interval, a probe grating appeared at the center of the screen. Its orientation either matched the last grating in the attended sequence or differed by ±4°. Participants reported whether the probe matched the final attended orientation by clicking the left mouse button for “Yes” and the right mouse button for “No.”

Each trial belonged to one of four counterbalanced conditions (attention to the left or right sequence, under either RTT or RDM structure). Participants completed five blocks of 36 trials each, with short breaks between blocks. The entire experimental session lasted approximately 50 minutes, not including EEG preparation.

### EEG acquisition and preprocessing

EEG data were recorded using a 128-channel BioSemi ActiveTwo system (Biosemi, Amsterdam, the Netherlands) with a sampling rate of 1024 Hz, and referenced online to the common mode sense (CMS) and driven right leg (DRL) electrodes. We additionally recorded two 2.5-minute resting state EEG with eyes opened and eyes closed, which were not used for the purpose of the analyses presented here.

EEG data were preprocessed using MATLAB (R2018b, The MathWorks, Natick, USA) and EEGLAB (v2021.1; Delorme & Makeig, 2004; Delorme et al., 2011). Data were downsampled to 250 Hz and low-pass filtered with a 40 Hz cutoff frequency. Each epoch was segmented from 500 ms before stimulus onset to 1 second after stimulus offset, resulting in epochs of 6.5 seconds in duration. Noise-contaminated channels and epochs were identified and removed following visual inspection.

Independent component analysis (ICA; Picard algorithm) was applied to identify and remove physiological artifacts such as muscle activity and eye blinks. Noise-related components were manually removed. Following ICA, “bad” electrodes were interpolated using the nearest-neighbor spline method, and the data were re-referenced to the average reference. Overall, 6.64% of electrodes were interpolated, while 1.67% of epochs and 7.33% of independent components were excluded.

### Behavioral Analysis

To investigate the RM effect induced by the rotating gratings, we analyzed the percentage of trials in which participants reported that the probe matched the last grating in the attended sequence. A two-way ANOVA was conducted with probe angle (same, +4°, or -4°) and the sequence type (RTT vs. RDM) as factors. To further assess the forward displacement effect, a post-hoc t-test was performed for the two conditions where the probe angle was +4°. Additional t-tests were conducted within each sequence type: for RTT, we compared +4° vs. 0° and +4° vs. −4°; for RDM, we compared 0° vs. +4° and −4° vs. +4°. These analyses aimed to confirm the presence of a forward bias in the RTT condition, but not in the RDM condition.

### SSVEP Analysis

The effects of SSVEP stimulation on the frequency content and topography of EEG activity were first assessed using a Fast Fourier Transform (FFT) applied to single-trial EEG data from 0 to 5 seconds following stimulus onset (both conditions combined; see Figure 1d-e).

To investigate the effects of attention and sequence type on SSVEP power, we conducted a time-frequency analysis using Morlet wavelet convolution with a Hanning taper. The analysis focused on the SSVEP frequency (5 Hz) and was restricted to the time window from 0.5 to 5 seconds to exclude transient EEG responses at stimulus onset. A sliding window of 1 second was used for the estimation.

To statistically compare conditions, we performed a cluster-based permutation test (cluster-level α = .05, α crit. = .05, two-tailed, 10’000 permutations) (Maris & Oostenveld, 2007). To assess the effects of attention, we contrasted SSVEP power between the attended hemifields (left vs. right) separately for the RTT and RDM conditions. The significant time window showing a bilateral effect of attention in the RDM condition (Figure 2b, dashed rectangle) was then used to generate scalp topographies of the attention effect in both conditions (Figure 2a-b).

To facilitate a direct comparison between conditions, we generated lateralization maps by flipping electrode coordinates so that the right hemisphere of the scalp consistently represented the contralateral side to the attended hemifield. These lateralization maps were then statistically compared between the RTT and RDM conditions, with cluster-based permutation statistics at the scalp level (Figure 2c).

### Population-Level Orientation Decoding

We applied the inverted encoding model (IEM; Brouwer & Heeger, 2009, 2011) to reconstruct population-level orientation representations from EEG time series. The encoding model consisted of 18 model basis functions, or “channels” (i.e., hypothetical neuronal populations), each centered on an orientation ranging from 0° to 170° in 10° steps. Individual channels were defined as half-wave rectified cosine functions raised to the power of 18.

For each trial, we constructed hypothetical channel outputs based on the stimulus orientation, resulting in a k × n matrix (C), where k = 18 (i.e., the number of orientation channels) and n is the number of trials. Each column in the C matrix represented the ideal response of the 18 channels to the stimulus orientation presented on that trial. The relationship between the observed EEG data and the hypothetical channel responses was modeled as:

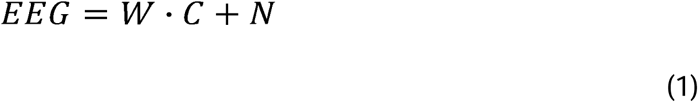

Where *EEG* is a matrix of size [*m* × *n*] with *m* electrodes and *n* trials, *C* is the hypothetical channel tuning function (CTF) matrix of size [*k* × *n*], representing tuning functions for *k* orientations across *n* trials, *W* is the weight matrix [*m* × *k*] mapping orientations to electrodes and *N* is the residual noise matrix.

To improve sensitivity to stimulus-selective neural responses, we selected a subset of 39 electrodes that exhibited the strongest responses at the SSVEP frequencies (responses exceeding the 75th percentile at 5 and 10 Hz; see SSVEP Analysis and Figure 1e).

Weights for each channel were derived using least squares estimation on a training dataset:

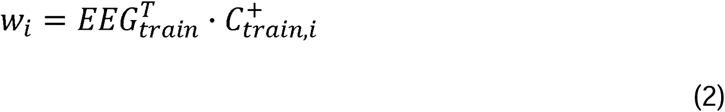

An optimal filter *V* was derived from the weights taking into account the noise correlation structure at each electrode, following the approach in (Kok et al., 2017). Using this approach, the CTF for the test dataset was reconstructed as:

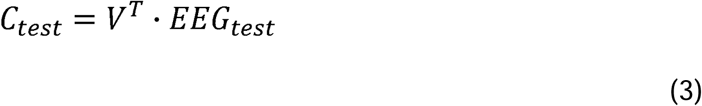

IEM was performed with a sliding time window of 40ms, using a leave-one-out cross-validation approach. The decoder was trained separately for each attended side and then the results were collapsed across conditions. Each trial was segmented into 25 shorter epochs, from 200 ms before the onset to 500 ms after the offset of each grating stimulus. This yielded approximately 1’100 epochs per participant per condition. To ensure equal sample sizes across participants, the first 1’000 epochs were selected and used for leave-one-out cross-validation. Estimated CTFs at each time point were circularly realigned so that the channel corresponding to the current stimulus orientation was centered at 0°, resulting in a [time points x 18 channels] matrix for each subject.

In the RTT condition, the orientations presented in consecutive segments were highly similar. As such, decoding of the current orientation could be influenced by ongoing EEG activity from the previous segment, which consistently included an orientation offset by -20° relative to the current segment. To mitigate this, we trained all decoders in the RDM condition to estimate the weights in eq. [2]. This approach also avoided systematic training biases due to varying EEG signal-to-noise ratios between conditions (Sprague et al., 2018).

To determine statistical significance, we employed nonparametric cluster-based permutation tests (Maris & Oostenveld, 2007) on the observed CTF metric. A surrogate distribution was generated by randomly shifting the CTF center before realignment (10’000 permutations). For both observed and surrogate CTF metrics, values exceeding the 95th percentile were identified, and neighboring significant pixels were grouped into clusters.

The sum of values within each cluster was recorded, and the maximum summed cluster values from surrogate data were used to form a null distribution. Observed clusters were considered statistically significant if their summed values exceeded the 95th percentile of this distribution (*p* < .05).

To quantify decoding performance, we computed a “decoding score” metric as the slope of a linear fit between the predicted CTF and a ground truth tuning function (i.e., a hypothetical CTF centered at 0°), yielding a time series of orientation decoding performance. A two-way repeated-measures ANOVA was performed on the decoding scores at each time point with factors attended side (left vs. right) and sequence type (RTT vs. RTT). To control for multiple comparisons, we performed time-resolved cluster-based permutation testing by shuffling condition labels and recomputing the ANOVA (10’000 permutations). Time points exceeding the significance threshold (uncorrected *p* < .05) were grouped into temporal clusters, and the maximum summed F-value in each permutation was stored to build a null distribution. Clusters in the observed data exceeding the 95th percentile of this null distribution were deemed significant (family-wise error rate: *p* < .05).

Biases in decoded CTFs were assessed by fitting a Gaussian function to the CTF averaged over significant time windows and estimating the deviation of the function’s center from 0°. A jackknife procedure (Miller et al., 1998) was employed to estimate the conditions, as well as from 0°. Specifically, the angular difference subsample, *D_-i_* was standard error of the angular difference between the centers of Gaussian fits across two calculated *N* times (where *N* is the number of participants), excluding one subject at a time. The standard error was then computed as:

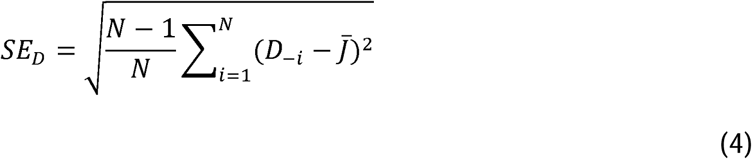

where *J̄* is the mean value of all subsamples, calculated as:

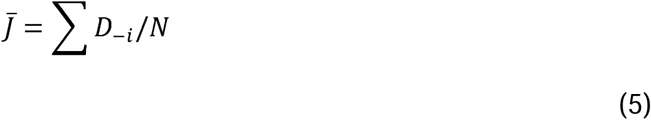

A jackknife t statistic, *t_j_*, was then calculated as

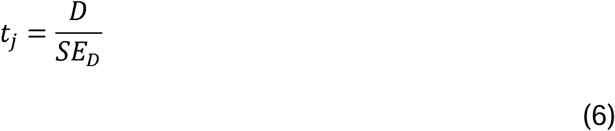

and the *p*-value was computed based on a degree of freedom of *N* - 1.

### Global Field Power (GFP)

Global field power (GFP) was calculated as the standard deviation of EEG potential across all electrodes at each time point within the decoding epoch (-200ms to 500ms relative to each grating onset). Higher GFP values indicate greater differences in activity across electrodes, reflecting stronger and more widespread neural activity.

To visualize the relationship between neural activity and orientation representation, GFP and the decoding score were averaged within the significant cluster identified in the decoding analysis (Figure 3a) for each subject. These values were then plotted in a 2D space. In Figure 4c, GFP is plotted after subtracting the subject-wise mean (i.e., the mean GFP across conditions for each subject) to facilitate visualization.

## Acknowledgements

We would like to thank Marius Peelen for comments and helpful discussions on the manuscript. This work was supported by the Swiss National Science Foundation (Grant numbers PZ00P1_179988 and PZ00P1_179988/2, and TMSGI1_218247).

## Data availability statement

Data will be made available in an online repository upon publication.

## Authors contribution

SB: Conceptualization, Data collection, Investigation, Writing – original draft.

MM: Conceptualization, Data collection, Investigation, Writing – review & editing.

DP: Conceptualization, Investigation, Writing – review & editing. Funding acquisition, Supervision.

